# An accurate and robust algorithm for solvent-excluded surface computation

**DOI:** 10.1101/294025

**Authors:** Lincong Wang

## Abstract

The solvent-excluded surface (SES^1^) of a molecule is an important geometrical property relevant to its solvation in aqueous solution and its interactions with other molecules. In this paper we present an accurate and robust analytic algorithm suitable for a large-scale SES analysis. The accuracy and robustness of our algorithm are made possible by an iterative strategy for the treatments of probe-probe intersections, probe-probe overlaps and limited numerical precision. The accuracy and robustness of our algorithm in turn make it possible to analyze on a large-scale the SESs for different types of proteins and various sets of ligand-protein interaction interfaces. The results illustrate the usefulness of SES and SES-defined physical and geometrical properties for the characterization of protein-solvent interaction and ligand-protein interaction where ligand could be either small molecule compound, or membrane lipid, or DNA or protein.

## 1 Introduction

The solvent-excluded surface (SES^1^) (also called molecular surface) of a molecule [1, 2] is a two-dimensional (2D) manifold that demarcates a topological boundary between the molecule represented as a finite union of atomic spheres with different radii (set 𝔸) and aqueous solvent modeled as an infinite union of probe spheres all with the same radius. A SES consists of three types of 2D boundary patches: convex spherical polygon (*∂*_*s*_(*i*)) on solvent-accessible atom i, saddle-shaped toroidal patch *(∂*_*t*_ (*i*, *j*)) determined by a pair ({*i, j*}) of accessible atoms and concave spherical patch *(∂*_*p*_(*i,j, k*)) by a triple ({*i, j, k*}) of accessible atoms. The set of solvent-accessible atoms 𝕊 is a subset of 𝔸 and each accessible atom is represented by a *∂*_*s*_. The *∂*_*p*_s are determined by a finite set ℙ of fixed probes while the *∂*_*t*_s are determined by both 𝕊 and ℙ. However in terms of probe only both *∂*_*s*_ and *∂*_*t*_ are defined by an infinite set of rolling probes. In contrast to SES the accessible solvent surface of a molecule [3, 4, 5] is formed only of *∂*_*s*_ polygons. Compared with either an accessible solvent surface or a van der Waals surface, a SES has finer details about the surface of a molecule since it also includes the areas determined by more than one atom and is thus expected to be better suited for the descriptions of protein-solvent interaction [6, 7, 8, 9, 10] and protein-ligand interaction [11].

At present more than ten SES computation algorithms [12, 2, 13, 14, 15, 16, 17, 18, 19, 20, 21, 22] have been published. Broadly-speaking they are either analytic or grid-based. In theory SES computation is straightforward if there is neither intersections nor overlaps among fixed probes and arbitrary-precision arithmetic could be used since then analytic equations [23, 5] exist for the exact computation of all the three patch types. In practice with limited numerical precision it is very challenging to handle algorithmically all the possible cases of probe-probe intersections and probe-probe overlaps. In addition the number of intersections or overlaps increases with molecule size. Of the five analytic SES programs [13, 14, 15, 16, 17] only PQMS and MSMS present some details for the treatment of probe-probe intersections. However, it seems neither PQMS nor MSMS could handle accurately and robustly all the cases of intersections or overlaps. When PQMS or MSMS detects error or inconsistency that it cannot correct, the program modifies atomic radius, typically by adding 0.1A to the radii of the atoms that generate the error, and then starts a new round of SES computation. For both PQMS and MSMS such modification and restart are frequent especially for large-sized biomolecules where several rounds of restart are routinely tried. Both PQMS and MSMS fail when the detected error could not be corrected with a fixed rounds of radius modification and restart. Such failures become frequent when the size of a molecule exceeds 10, 000 atoms. The arbitrary modification to radius and restart introduce inconsistency, reduce the precision of the computed SES area and make them less suitable for a large-scale SES analysis. A possible advantage of grid-based algorithms [19, 20, 21, 22] is that their computed surfaces are less sensitive to either probe-probe intersections or probe-probe overlaps or limited numerical precision especially when a large grid is used. However these algorithms increase cubically with grid resolution in time and in space. In addition they could not properly assign area to each individual surface atom and the computed total area has limited precision especially for large-sized molecules. The main goal of our analytic and probe-centric algorithm is to compute as accurately and robustly as possible the SES area for each patch. The goal is achieved through (1) an iterative strategy for 𝕊 and ℙ computation that minimizes the impact of limited numerical precision, (2) the separation of probe-probe interactions into three different types, and (3) the accurate and robust treatments of all the cases of probe-probe intersections and probe-probe overlaps with no modifications to atomic radii, no perturbations to atomic positions and no program restart. As shown in our companion papers [24, 25], the accuracy and robustness of our algorithm make it possible to analyze on a large-scale SES and SES-defined physical and geometrical properties for different types of proteins and ligand-protein interaction interfaces. The results of our large-scale analyses illustrate the important roles played by SES area and a list of SES-defined properties in protein-solvent interaction and ligand-protein interaction where ligand could be either small molecule compound, or nucleic acid, or membrane lipid or protein.

## 2 The geometry of SES

The solvent-excluded surface (SES) of a molecule is composed of three types of 2D patches: (1) convex spherical polygons (*∂_s_*(*i*)s) on a set 𝕊 of accessible atoms, (2) saddle toroidal patches (*∂*_*t*_(*i, j*)s) defined by atom pairs ({*i,j*}s), and (3) concave spherical polygons *(∂_p_(i,j,* k)s) on a set ℙ of fixed probes (**p***ijk*^s^) whose positions are determined by atom triples ({*i, j, k*}s) (Fig. 1). There exists the following relationship among the three types: the polygon *∂*_*s*_(*i*) on atom i is surrounded by an array of interleaved *∂_t_(i,j*)s and *∂*_*p*_(*i, j, k*)s^2^. The *∂*_*t*_(*i,j*)*s* in the array are traced out by a rolling probe that starts at the first fixed probe p_*ijn*_*j*_1_ and then rolls along the major circle of a torus determined by atom pair {*i, j*1} until it hits atom *j*_2_ where a spherical polygon *∂*_*p*_(*i, j_1_, j*_2_) appears, that is, the rolling probe reaches the second fixed probe p_*i*_*j*_1*j*2_. The probe then rolls along the major circle of a torus determined by atom pair {*i, j*_2_}, and continues rolling until it reaches the last fixed probe p_*i*_*j*_n-1_*j*_n_, then rolls along the last atom pair {*i*, *j*_*n*_} and finally stops at the last atom *j_n_.* This relationship is the geometrical foundation for analytic SES computation.

**Figure 1:**
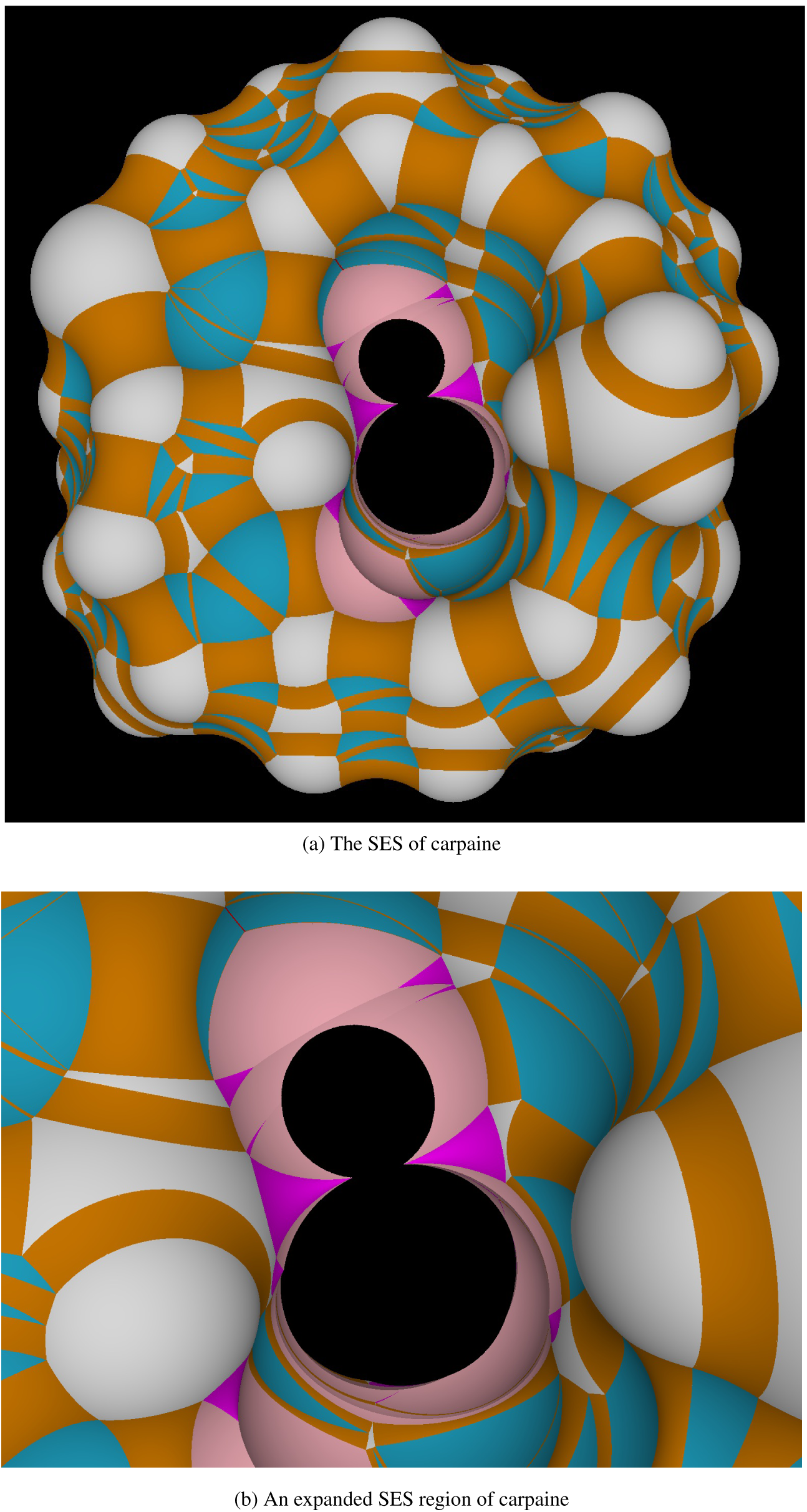
The three types of SES patches on carpaine, a papaya alkaloid. **(a)** The SES of carpaine viewed from the top. The spherical polygons (*∂*_*s*_(*i*)s) of accessible atoms are colored in white, the toroidal patches (*∂*_*t*_(*i, j*)s) in orange for regular torus and in magenta for spindle torus, and the spherical polygons (*∂*_*p*_(*i,j, k*)s) on fixed probes in cyan for isolated probes and in pink for intersected ones. **(b)** An expanded region composed mainly of intersected *∂*_*p*_(*i, j, k*)s and spindle toroidal patches. All the figures in this paper are prepared using our structural analysis and visualization program written in C++/Qt/OpenGL. The triangulated SES is rendered using Phong shading model with no specular lighting.

The above SES geometry shows that a simple algorithm could be developed if a *∂*_*t*_ (*i, j*) is followed by a *single ∂_p_(i,j, k)* and a single *∂*_*p*_(*i, j, k*) is followed by another single *∂*_*t*_(*i, k*) and so on. No much complications arise if a fixed probe in the array intersects only with its consecutive fixed probes. However the surface of a protein especially a large-sized protein could have geometrically complicated regions such as ligand binding sites where a *∂*_*t*_(*i, j*) is followed not by one, but a *set* k of third atoms (Figs. 1b and 2). There will be a set P*_i_j* (k) of fixed probes each of them determined by a *k* ϵ k and the same atom pair {*i, j*}. One challenge of SES computation is that there may exist complicated intersections among the fixed probes in P *ij* (k) (Fig. 2). Furthermore, two fixed probes may intersect with each other without sharing any atom and thus their intersection could not be discovered by tracing a probe rolling only between individual pairs of atoms. The existence of complicated probe-probe intersections is often cited as an undesirable feature of SES when compared with accessible solvent surface. How to treat probe-probe intersection determines largely the accuracy and robustness of any analytic SES computation algorithm. In theory P_*ij*_ (k) could be discovered by carefully tracing a probe that rolls around atom i as having been implemented in previous analytic SES programs [13, 14, 15]. In practice, however, the exhaustive treatments of all the possible cases of probe-probe intersections and overlaps are greatly exacerbated by limited numerical precision.

**Figure 2:**
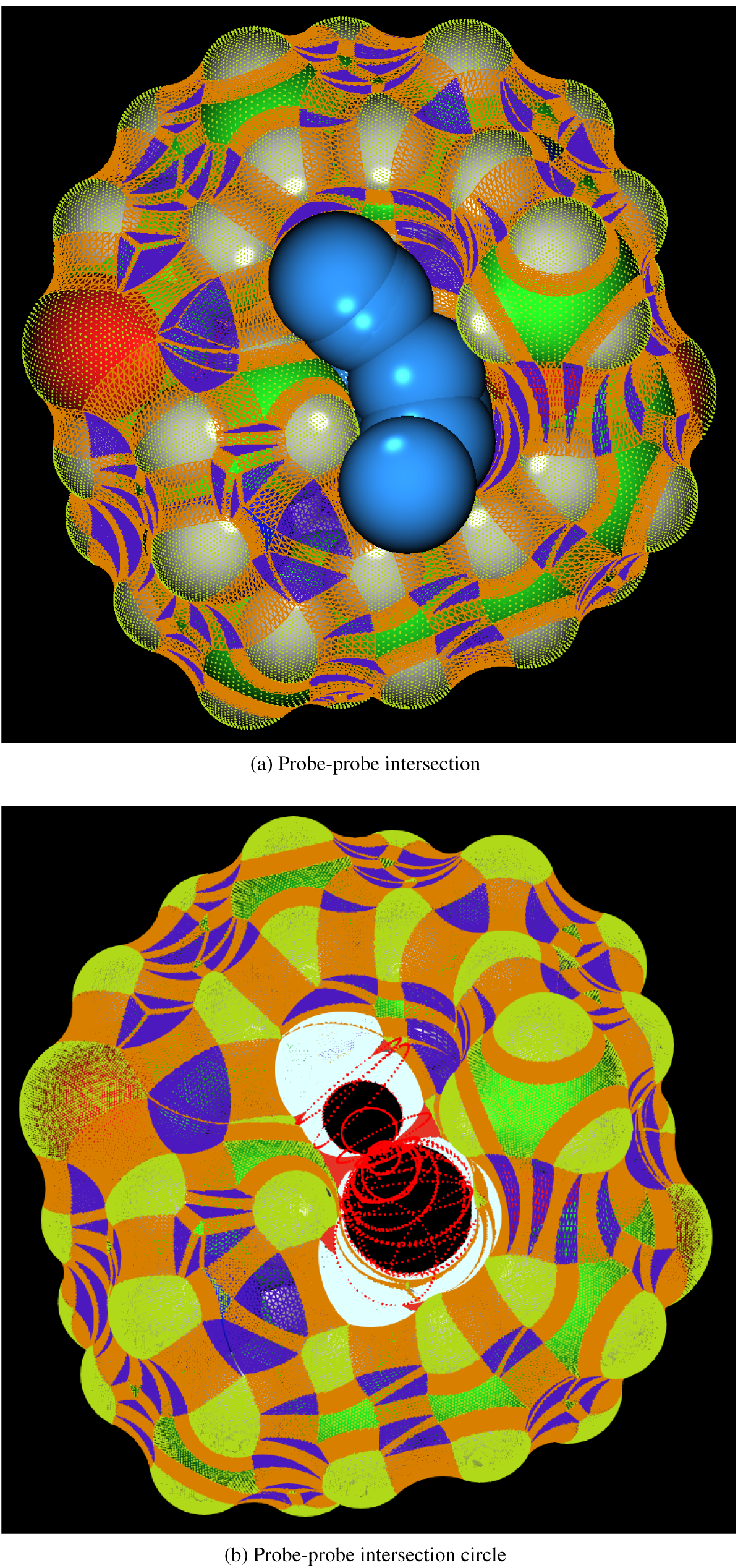
Probe-probe intersections and intersection circles of carpaine. **(a)** A set of intersected fixed probes colored in azure. **(b)** Probe-probe intersection circles 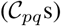 colored in red. In both **(a)** and **(b)** convex spherical polygons are rendered as points and colored in acid yellow, toroidal patches as wire-frames and colored in orange for regular torus and in red for spindle ones, isolated concave spherical polygons as wire-frames and colored in blue while intersected concave spherical polygons as points and colored in white. The molecule itself is depicted in a space-filled model.

## 3 The algorithm

Unlike previous analytic SES computation algorithms [13, 14, 15] we postpone the treatments of probe-probe intersections and overlaps to a late stage when all the accessible atoms and all the fixed probes have been discovered. Knowing all the fixed probes makes it easier to first detect all the possible cases of probe-probe intersections and probe-probe overlaps and then treat them accurately and robustly. An additional advantage of having all the fixed probes is that it becomes possible for our algorithm to neither increase atomic radii as required in both PQMS [13] and MSMS [14] nor perturb atomic positions as required in ⍺-shape [26]. Our algorithm and its implementation are somewhat complicated due mainly to the requirement to detect and treat various probe-probe interactions and overlaps, to deal with limited numerical precision and to *visually* verify the computed SESs and SES areas. In the following we present the key steps of our algorithm with a brief mention of the other steps.

### 3.1 The computation of accessible atoms and fixed probes

A prerequisite for accurate and robust analytic SES algorithm is the computation of (1) the set 𝕊 of all the accessible atoms on both the exterior surface of a molecule and the surfaces of its interior cavities and (2) the set ℙ of all the fixed probes. Though analytic equations exist for their computation [23], to minimize the impact of limited numerical precision our algorithm employes an iterative strategy. An atom range search tree **R**_*a*_ is first constructed for all the atoms and used to speed up the computation of the SA-adjacent list l_*s*_(*i*) for accessible atom *i.* Two atoms *i, j* are SA-adjacent if *d_ij_ < r_i_ +* 2*R*_*p*_ + *r*_*j*_ where *d*_*ij*_ is the distance between atom i and atom *j*, *r*_*i*_, *r*_*j*_ their respective radii and *R*_*p*_ the probe radius. The sides of *∂*_*s*_(*i*) are determined by the intersections between atom i and the individual atoms in l_*s*_ (*i*).

The algorithm starts with the computation of set 𝕊, area *a*_*s*_ and SA-adjacent list l_*s*_ for each accessible atom.

1. CONSTRUCT atom range search tree R_*a*_
2. COMPUTE set 𝕊
3. FOR each atom *i* ϵ 𝕊
4. COMPUTE area *a*_*s*_ (*i*) and adjacent list l_*s*_ (*i*)

The accessible area of atom i is computed as, *a*_*s*_(*i*) = *n*(*i*)*a*_*min*_, where *n*(*i*) is the number of vertices on atomic sphere i that are not buried by any other atoms and *a*_*min*_ the area assigned to a single vertex. In our implementation, each atom is represented as a sphere and an atomic sphere is in turn represented by a set of uniformaly-distributed vertices. A similar vertex representation is used for each probe sphere. An atom is added to 𝕊 if it has at least one unburied vertex. The list l_*s*_(*i*) for atom i is found using R_*a*_ and may include atoms that do not belong to 𝕊. An atom in l_*s*_ (*i*) that is not in 𝕊 must have *a*_*s*_ < *a*_*min*_.

#### The set of fixed probes ℙ is computed as follows.

1. LET set Q = 𝕊_p_ = ∅
2. FOR each atom *i* ϵ 𝕊
3. FOR each atom pair {*J, K*} in l_*s*_ (*I*)
4. IF atom triple {*I, J, K*} ∉ Q
5. ADD {*i, j, k*} to Q
6. COMPUTE fixed probe p _*ijk*_
7. IF **P**_*ijk*_ is a valid probe
8. ADD **P** _*i j k*_ to ℙ
9. IF *j* ∉ 𝕊
10. ADD *j, k* to 𝕊_p_ and LET *a*_s_ (*k*) = *a*_*mm*_
11. FOR each atom *j* ϵ 𝕊_p_
12. IF *j* ∉ 𝕊
13. ADD atom *j* to 𝕊
14. CONSTRUCT probe range search tree R_*P*_

A fixed probe that does not collide with any protein atom is called a *valid* probe. Up to two valid probes may exist for the same atom triple. For any triple {*i, j, k*} though *i ϵ* 𝕊 either *j* or *k* may not belong to 𝕊. Additional valid probes if exist are first computed using all the triples in 𝕊_P_ and then added to ℙ (step 8) while each atom of such a triple that produces a valid probe is added first to 𝕊_P_ (step 10) and then to 𝕊 (step 11-13). Finally a probe range search tree R_*p*_ for ℙ is constructed and used late to expedite probe-probe intersection computation. The three contact points, v_*i*_, V_*j*_ and v_*k*_, between a fixed probe p_*ijk*_ and atom triple {*i,j, k*}, define respectively a spherical triangle Δ_*ijk*_ on the probe sphere and three great circles, 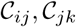 and 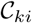, that pass respectively through its three sides (Fig. 2b). Let o_*p*_ be probe center and **n** _*ij,*_ **n**_*jk*_, **n**_*ki*_ three normals that are perpendicular respectively to the three plane triangles Δ_*oij*_, Δ*ojk* and Δ_*oki*_, then the set (**V**_*ijk*_(**p**_*ijk*_)) of the vertices on probe sphere \ that are inside spherical triangle Δ_*ijk*_ could be determined by checking their dot products with **n**_*ij*_, **n**_*j k*_ and **n**_*ki*_.

### 3.2 The detection of probe-probe interactions

The next step is the detection of all the possible interactions among the set of fixed probes ℙ. Two fixed probes interact with each other if they either *share* at least one atom, or *overlap,* or *intersect* with each other. Two fixed probes could share up to 3 atoms. It is important to separate atom-sharing from probe-intersecting since the latter may exist without sharing any atom. This step begins with the detection of probe-probe interactions and then separates them into three types: repetition, intersection and atom-sharing.

1. FOR each probe p ϵ ℙ
2. FOR each probe **q** ϵ **l**_*P*_(**p**)
3. **IF** *d*_pq_ < D_P_,min
4. ADD probe q to set **Q**_O_ (**P**) {probe-probe overlap}
5. ELSE IF *d*_pq_ < d_*p,max*_
6. ADD probe q to set **Q**_I_ (**P**) {probe-probe intersection}
7. ELSE IF p and **q** share only two atoms *i* and *J* {atom-sharing}
8. INSERT atom pair {*I, J*} as a key and probe pair {p, q} as a value into map m_t_
9. ELSE if p and **q** share **3** atoms *I,J,K* {atom-sharing}
10. INSERT each of the three pairs ({*i, j*}, {*j, k*}, {*i, k*}) as a key and probe pair {**p, q**} as a value into **M**_T_

where **l**_*p*_(**p**) is the adjacent list for probe p computed using probe-probe range search tree **R**_*p*_, and *d*_*pq*_ is the distance between probe p and probe **q**. Threshold *d_p_,*_*min*_ is used to determine whether two probes overlap with each other and *d_p_,_max_ = 2R_p_.* Sets **Q**_O_ (**p**) and **Q**_I_ (**p**) include respectively all the fixed probes that repeat (overlap) with probe p and intersect with probe p. A key in map **M**_T_ is an atom pair while the key’s value is an array of fixed probes that share the same atom pair. A probe may repeat or intersect with several other fixed probes. Probe repetition if ignored will lead to the same area *a*_*p*_ being computed more than once. In our algorithm a subset of unique probes are selected from **Q**_O_ for SES computation. Map **M**_T_ is similar to the reduced surface of MSMS [**14**] but in our algorithm it is used only for toroidal patch computation rather than for the discovery of fixed probes as in MSMS.

### 3.3 The computation of toroidal patches

Each item in map defines a toroidal patch. Specifically the key of an item, an atom pair {*i, j*}, defines a major circle of radius *R*_*t*_ while its value, a set of fixed probes, determines the starting and ending positions for a set of arcs 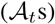 on the major circle along them toroidal patches are traced out by a rolling probe. The minor circles of all the tori have the same radius as each other: that is, the probe radius *R*_*p*_. There exist two types of torus: regular and spindle (Fig. 1 colored respectively in orange and magenta). They could be separated from each other easily at least in theory: if *R*_*t*_ > *R*_*p*_ it is a regular torus, otherwise a spindle torus. The computation of the arc 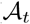 and toroidal patch for key {*i, j*} is straightforward if the key’s value is a single pair of probes. However, if the key’s value is a set of fixed probes, these probes need to be ordered first along the major circle and there may exist more than one toroidal patch for an atom pair.

1. LET set 𝕊_T_ = ∅
2. FOR each key (atom pair {*i, j*}) in **M**_T_
3. COMPUTE *R*_*t*_
4. IF *R*_*t*_ < *R*_*P*_
5. MOVE the item from **M**_T_ to M_ST_
6. ELSE
7. COMPUTE regular toroidal patch *∂*_*t*_ (*I, J*)
8. COMPUTE AREA *A*_*T*_
9. IF *i, j ∉*𝕊
10. ADD {*i, j}* to 𝕊_T_ and set *a*_*s*_(*I*) = *a*_*s*_(*J*) = *a*_*min*_
11. FOR each atom *j ϵ* 𝕊_T_
12. IF *j ∉* 𝕊
13. ADD ATOM *j* to 𝕊

where **M**_ST_ is a map for spindle torus. This step computes only regular torus. If atom *i* or *j* does not belong to 𝕊 it is added to 𝕊_T_. If 𝕊_T_ ≠ ∅ additional valid probes if exist are first computed using all the atom triples in ℙ and then appended to P while each atom of such a triple that produces a valid probe is added to 𝕊 (step 11-13). The newly-added probes are checked for their interactions among themselves and with the previous ones. A probe pair in M_ST_ also belongs to a set of intersecting probes Q_j_. The area *a*_*t*_ of a regular toroidal patch *∂*_*t*_ is computed analytically.

A spindle toroidal patch is computed similarly as a regular torus. The main difference is that for a spindle torus the two fixed probes that define the starting and ending positions of a rolling probe intersect with each other at two positions **v**_*i*_ and **V**_*j*_.

1. FOR each key (atom pair {*i, j*}) in **M**_ST_
2. COMPUTE great circle *c*_*ij*_
3. COMPUTE the intersecting circle *c*_*pq*_ between probe p and probe q (the key’S value)
4. IF *c*_*ij*_ and *c*_*pq*_ intersect
5. COMPUTE their intersecting points v_*i*_ and v_*j*_
6. COMPUTE areas *a*_*t*_(*i*) and a_t_(*j*)
7. ELSE
8. COMPUTE *∂*_*t*_ as a regular torus

The great circle 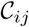 passes through side {*_i, j_*} of spherical triangle Δ_*ijk*_ on probe **p**_*ijk*_. Spindle patch area *a*_*t*_(*i*) is computed via triangulation as detailed in visual verification and algorithm implementation section. Due to limited numerical precision it is possible that 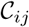 and 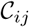 does not intersect but atom pair {*i, j*} is in **M**_ST_.

### 3.4 Probe-probe intersections

The most difficult step in SES computation and triangulation is used to be the determination of all the possible intersections among fixed probes. In PMQS, MSMS and contour-buildup [15] algorithms probe-probe intersections are dealt with when they are discovered greedily via atom-sharing. However two probes may intersect with each other even they may not share any atom. Such an intersection could not be discovered through atom-sharing. The exclusion of such intersections leads to errors in SES area and defects in triangulated surface. Unlike them our algorithm first computes all the valid fixed probes and then detect exhaustively all the possible probe-probe interactions. We believe such an approach could best handle various cases of interactions originated from either complicated local geometry or limited numerical precision or both. Specifically in our algorithm both S and P have been computed at this step and consequently we have essentially an adjacent list representation for a probe-probe interaction graph. It is straightforward to remove all the vertices in ***V***_*ijk*_ (**p**_*ijk*_) that are intersected by any other fixed probes.

1. FOR each probe **P**_*IJK*_ Ȉ ℙ
2. IF Q_I_ (**P_*ijk*_**) ≠ ∅
3. FOR each probe q Ȉ q_j_ (p_*ijk*_)
4. REMOVE all the vertices in v_*ijk*_ (p_ijk_) that are buried by q
5. COMPUTE AREA *A_P_(I,J,K*)

where {*i, j, k*} is the atom triple that defines probe **p**_*ijk,*_ and **V**_*ijk*_ (**p**_*ijk*_) is a set of vertices that are inside spherical triangle Δ_*ijk*_ on probe sphere **p**_*ijk*_. The area *a*_*p*_(*i,j,k*) of a concave spherical polygon *∂*_*p*_ (*i,j,k*) of an intersected probe (colored in pink in Fig. 1) is computed as, *a*_*p*_(*i,j, k*) *= n*(*i,j, k*)*a_min_,* where *a*_*min*_ is the area assigned to a single vertex on the probe sphere. A fixed probe that does not intersect with any probe is called an *isolated* probe (colored in cyan in Fig. 1). The area of an isolated probe is computed analytically.

## 4 Visual verification and algorithm implementation

The correct treatments of all the possible cases of probe-probe interactions and limited numerical precision pose major challenges for accurate and robust SES computation and triangulation. Particularly due to limited numerical precision it is difficult to guarantee that the computed SES is accurate for a region where there exist complicated probe-probe interactions (Figs. 1b, 2 and Fig. S1 of the Supplementary Materials). Our experience shows that one way to detect SES computation error is to follow the paradigm of what you see is what you get by rendering individual SES patches as geometric primitives. Furthermore to detect SES computation error it is necessary to visualize various intermediate geometries such as the initial spherical triangle (*∂_p_*) on a fixed probe, the major circle and arc of either a regular torus or a spindle torus, the intersecting circle 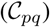 between two fixed probe spheres (Fig. 2b), and the intersecting points (**v**_*i*_s and *V*_*j*_s) between a probe great circle and an intersecting curve 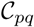 (Fig. 2b and Fig. S1). For the purpose of error detection via visual inspection most of these intermediate geometries are rendered as either points or lines or wire-frames. For examples, both regular and spindle toroidal patches are triangulated and rendered as wire-frames with and without shading (Fig. 2 and Fig. S1). The concave spherical triangle (*∂*_*p*_) on an isolated fixed probe is triangulated via subdivision and rendered similarly. Both the *∂*_*p*_s on pairs of intersecting probes and the convex spherical polygons (*∂*_*s*_s) on individual accessible atoms are rendered as points. When an error in SES is detected visually, the responsible accessible atoms and fixed probes as well as the relevant intermediate geometries are selected, labeled and analyzed to assess whether the error is caused by limited numerical precision or represents a previously-untreated case of probe-probe interaction or both. Through the visual inspection of more than 200 proteins, most of them are large-sized ones with more than 20,000 atoms, we are able to identify various errors and to further develop accurate and robust remedies for them. The number of errors in SES and the number of defects in triangulated surface increase with molecule size. In fact, our iterative strategy for the computation of both 𝕊 and ℙ was developed in order to eliminate the various errors detected through visual inspection. As illustrated in Fig. S1 of the Supplementary Materials finding errors through such a process is definitely time-consuming and tedious. The pay-off is that we are able to treat accurately and robustly various probe-probe interactions with no need to modify atomic radii or to perturb atomic positions even for large-sized proteins.

In our implementation both accessible atoms and fixed probes are represented by a set of uniformly-distributed 40, 962 vertices for molecules with ≤ 40, 000 atoms. For molecules with > 40, 000 atoms they are represented by 10, 242 vertices. The atomic radii used in our implementation are as follows: C=1.70Å, N=1.55Å, O=1.52Å, S=1.75Å, H=1.09Å and Se=1.80Å. The probe radius *R*_*p*_ is set to 1.4&#x00C5;. The threshold *d_p_,*_*min*_ used in the detection of probe-probe overlapping is set to 1.0 × 10 ^6^ Å.

## 5 Results and discussion

The main goal of our algorithm is to compute the SES areas of both individual patches and individual accessible atoms as accurately and robustly as possible rather than to achieve the best visual effects for molecular surface [27]. In the following we first describe the key differences between our algorithm and previous analytic SES algorithms. We then discuss the precision of the computed SES area as well as the robust and accurate treatment of probe-probe interaction. Finally we compare the SESs for several molecules by our algorithm and **MSMS**. Of the five analytic SES computation programs [13, 14, 15, 16, 17] only **MSMS** is currently available for free download.

### 5.1 The differences between previous analytic SES algorithms and our algorithm

Our algorithm relies on the same set of analytic equations as previously-derived for SES patch computation [23] and thus has overall resemblance with published analytic SES computation algorithms [13, 14, 15, 16, 17]. However, our algorithm is probe-centric and thus differs from either torus-centric **PQMS** [13] and **MSMS** [14] or atom-centric contour-buildup algorithm [15]. The other key differences include (1) an iterative strategy for 𝕊 and ℙ computation to minimize the impact of limited numerical precision, (2) the separation of probe-probe interactions into three different types, (3) the accurate and robust treatment of all the cases of probe-probe intersections, and (4) no modification to atomic radii, no perturbation to atomic positions and no program restart. With no modification to atomic radii and no perturbation to atomic positions, the SES area computed by our algorithm should be more accurate than those by both **PQMS** and **MSMS** that modify atomic radii and that by **ALPHA**S**HAPE** [16] that perturbs atomic positions. In contrast with our algorithm **MSMS** requires manual intervention for the computation of the SES for any interior cavity. As far as application is concerned the frequent failure of **MSMS** for molecules with > 10, 000 atoms makes it ill-suited for large-scale analyses. In contrast, our SES program rarely fails even for very large-sized proteins as long as the PDB file itself has been properly processed.

### 5.2 The precision of SES patch areas

In our algorithm both the SES patch area (*a*_*p*_) of a spherical triangle on an isolated fixed probe and the area (*a*_*t*_) of a regular toroidal patch are computed analytically. However the patch area (*a*_*s*_) of a spherical polygon on an accessible atom, the patch area (*a*_*p*_) of a spherical polygon on an intersected fixed probe, and the area of a spindle toroidal patch are computed approximately. We do not compute *a*_*s*_s analytically since they could be estimated with high precision during 𝕊 computation. The analytic equations for the computation of the *a*_*p*_ for a highly-intersected fixed probe (Fig. 2) though exist are rather complicated. The precision of the approximately-computed SES areas could be estimated as follows. For a molecule with ≤ 40,000 atoms each of its accessible atoms or each of its fixed probes is represented by 40,962 vertices or 81, 920 triangles. For a probe with *R_p_ =* 1.4Å, each vertex represents an area of *a_min_ =* 6.013 × 10&3×2014; and each triangle has an area of 3.0066 × 10^−4Å2^. If these triangles are converted into a grid, the corresponding grid resolution is about 2.5 × 10 ^2Å^. For a molecule with > 40, 000 atoms each of its accessible atoms or each of its fixed probes is represented by 10, 242 vertices or 20, 480 triangles. In our implementation, a sulphur atom has the largest radius, 1.75Å, and thus its *a*_*s*_ has the largest uncertainty. For a sulphur atom each vertex represents an area of *a*_*min*_ = 3.758 × 10^−3^Å^2^ and each triangle has an area of 1.879 × 10^−3^Å^2^. If these triangles are converted into a grid, the corresponding grid resolution is about 6.6 × 10^−2Å^. Å grid resolution of 6.6 × 10^−2^Å is still a magnitude smaller than a typical resolution used in a grid-based method even for a relatively small molecule with about 5, 000 atoms. The area *a*_*t*_ of a spindle toroidal patch is computed via triangulation with a grid-resolution of 1.2°. The area of a plane triangle for a toroidal patch differs from its exact value by a constant amount &#x03B4;_a_ that is at most 1.5 × 10^−5^Å^2^. The number of plane triangles whose areas may deviate from their exact values by &#x03B4;_*a*_ is estimated to be < 1, 000 for the largest spindle tori. Thus the area of a spindle toroidal patch is at most 1.5 × 10^−2^Å^2^ smaller than its exact value. Most toroidal patches have < 200 plane triangles whose areas may deviate from their exact values by &#x03B4;*a* and thus the corresponding errors in the computed *a*_*t*_s should be < 3.0 × 10^−3^ Å^2^. In summary in the current implementation of our algorithm the error for each patch is estimated to be < 0.01Å^2^.

### 5.3 Accurate and robust treatment of probe-probe interactions

Though it is important to be able to compute with high precision the SES areas of both individual patches and individual accessible atoms, a more challenging problem for SES computation is the treatments of all the cases of probe-probe interactions and limited numerical precision. Untreated cases of probe-probe intersections or probe-probe overlaps will likely lead to errors in the computed areas and defects in the triangulated surfaces (the Supplementary Materials Figs. S2 and S5). In fact the poor treatment of probe-probe interactions may also lead to defects in a triangulated surface generated by a grid-based approach (Fig. S2a). In our algorithm the treatment of probe-probe interactions is postponed to a late stage when all the accessible atoms and all the fixed probes have been computed. By means of visual inspection and verification we find that it is easier to design methods to deal with probe-probe interactions when all the fixed probes are known though from a pure algorithmic viewpoint it is desirable to treat them as early as possible. The accurate and robust treatment of all the cases of probe-probe interactions and the high precision of the computed patch area are two most salient features of our algorithm. Ås shown in our companion papers [24, 25] these features make our algorithm ideally suited for large-scale applications to various types of proteins and ligand-protein complexes.

### 5.4 Comparisons with msms and grid-based algorithms

Åt present more than ten SES programs of either analytic or grid-based have been published [13, 14, 15, 16, 28, 17, 18, 19, 20, 22]. &#x00C5;mong the five analytic programs only MSMS is currently available for free download. In the following we first describe the differences between the triangulated surfaces generated by MSMS and our algorithm including a discussion about the difficulty to compare the two programs patch-by-patch or atom-by-atom. We then use the molecular surfaces of carpaine (Fig. 3) to illustrate the differences between our algorithm and two grid-based programs.

**Figure 3:**
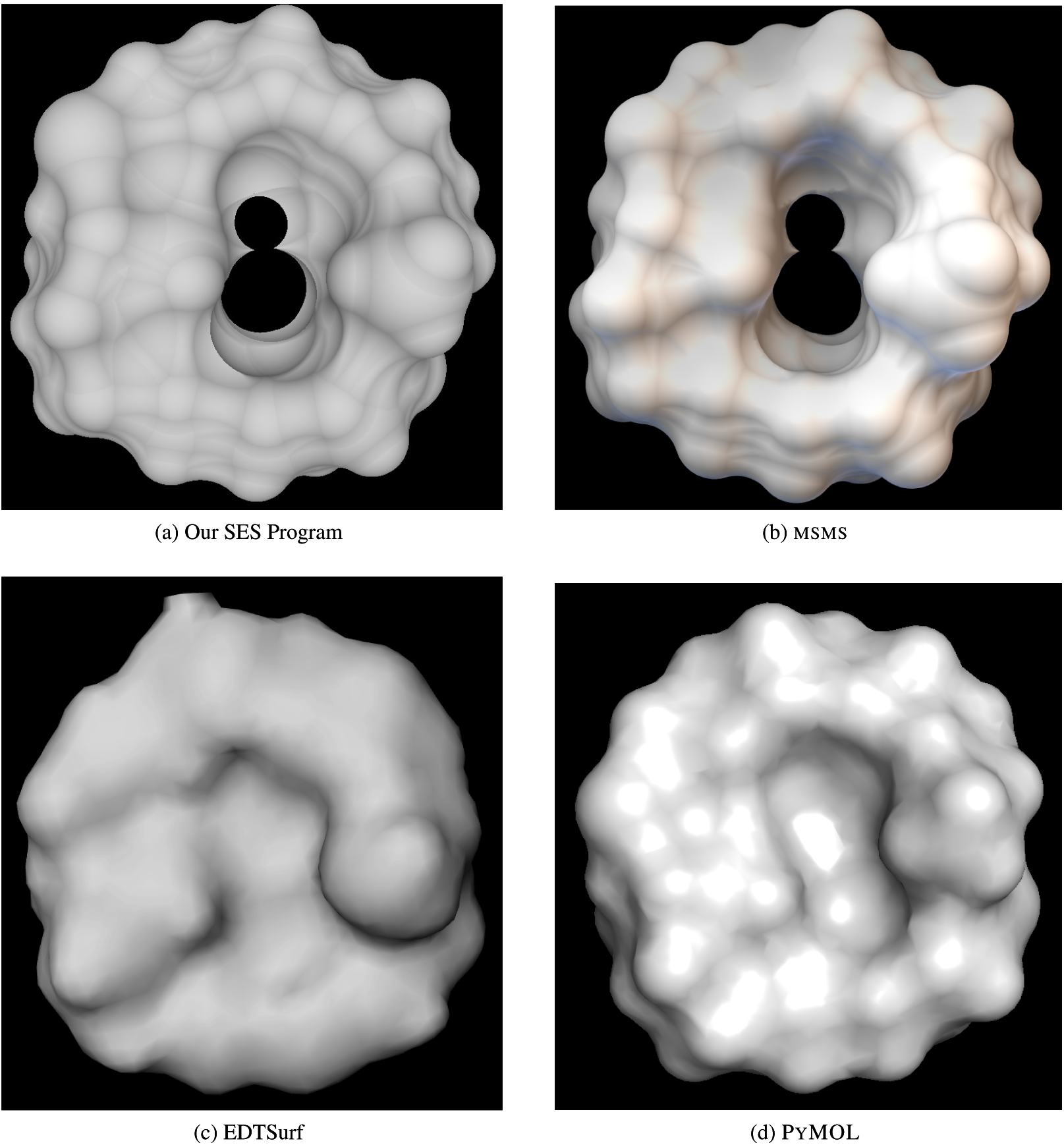
The surfaces of carpaine by four programs. **(a)** The SES by our algorithm is computed using 40, 962 vertices for sphere representation. **(b)** The MSMS surface is generated using a *R_p_ =* 1.2Å, density=100 and high density=100. **(c)** The molecular surface by EDTSurf is generated using IVIEW [30]. **(d) Å**s a reference the molecular surface generated by a popular visualization program PyMOL lacks fine details especially for intersected regions. The middle hole in the molecular surface by either EDTSurf or PyMOL is filled.

The MSMS SESs used for comparison are generated by PMV [29], a program included in MGLTools (version 1.5.6). The probe radius *R*_*p*_ for MSMS surfaces is set to either 1.2Å or 1.3Å since our comparisons show that the MSMS surfaces generated by these two radii are most similar to those by our algorithm with *R*_*p*_ = 1.4Å likely because MSMS uses atomic radii larger than ours. It is difficult to compare the areas of individual patches or individual atoms because they are calculated using different atomic radii and because probe-probe interactions are treated differently. A patch-by-patch comparison is further complicated by the fact that **MSMS** almost always increases atomic radii for some atoms, and either clashes or responds very slowly to user input when the resolution for triangulation is high and the molecule is large. Though it is impossible for our algorithm to generate a surface triangulation that exactly matches that by **MSMS** and though no details were given in their paper, **MSMS** triangulation is supposed to follow the same analytic equations as those used for area computation. As described above in our algorithm the same set of triangles are used for both the area computations and triangulations of spindle tori and the spherical polygons on intersected probes. For comparison with **MSMS** an isolated probe and an accessible atomic sphere are both triangulated with 2, 560 triangles while a regular torus is triangulated with a grid-resolution of 1.2°. As shown in Figs. 3a and 3b and Figs. S3-S5 of the Supplementary Materials the triangulated surfaces generated by our algorithm have close overall resemblance with those by **MSMS** except for the regions with complicated probe-probe interactions. Specifically the **MSMS** SESs lack the sharp edges that demarcate different concave spherical polygons on intersected probes. However we are not sure whether the lack of such sharp edges affects the precision of the computed SES areas since **MSMS** presents no details for the area computation of a highly-intersected region.

A direct comparison between a grid-based algorithm and our analytic SES algorithm is tricky since the former does not assign surface area to individual patches and atoms. In addition it is rare for a grid-based algorithm to use a resolution of < 0.5&#x00C5; for a molecule with > 40, 000 atoms. As described in section 5.2, if converted into grid-resolution, our implementation has a resolution of < 6.6 × 10^−2&^#x00C5; even for large-sized molecules. The superior accuracy of our algorithm is easily visible when we compare the SES of carpaine by our algorithm with the molecular surfaces generated by two grid-based methods: EDTSurf [22] and PyMOL. As shown in Fig. 3 in the middle of carpaine lies a hole where complicated probe-probe interactions occur (Fig. 2b). The surfaces generated by the two grid-based methods have the hole covered possibly because they have used atomic radii larger than ours or their grid-resolutions are too large to have such fine details. Compared with the analytic SES by either our algorithm or **MSMS** the surface by either EDTSurf or PyMOL is rather coarse.

## 6 Conclusion

A robust, probe-centric and analytic algorithm has been developed for the accurate computation of solvent-excluded surface (SES). Two salient features of our algorithm are (1) the exhaustive treatments of various cases of probe-probe interactions and proper handling of limited numerical precision through an iterative strategy for the computation of both solvent-accessible atoms and fixed probes, and (2) accurate SES computation with no modifications to atomic radii and no perturbations to atomic positions. Large-scale applications to various types of proteins and ligand-protein interaction interfaces confirm the accuracy and robustness of our algorithm and support the importance of a list of SES-defined physical and geometrical properties to protein-solvent interaction and protein function.

## Supplementary Materials

## S1: Visual inspection and verification

The complexity of probe-probe interactions in combination with limited numerical precision makes it necessary to visually verify both the computed SES areas and the triangulated SESs. As illustrated in Fig. S1 an analytic SES algorithm may fail to compute a few patches even it could correctly compute tens of thousands of other patches for the same molecule. During the development stage we have detected many similar defects through visualization and have removed them algorithmically.

### S2: The defects in triangulation by a grid-based algorithm

As with an analytic SES algorithm the poor treatment of probe-probe intersections by a grid-based algorithm may also lead to defects in the generated surface (Fig. S2). PyMOL is a widely-used molecular visualization program with molecular surface computed using a grid-based algorithm.

### S3: The comparisons of the SESs by our SES algorithm and MSMS

In this section, we compare the SESs for three proteins with increasing sizes by our program and MSMS. The MSMS surfaces are generated by PMV, a program included in MGLTools (version 1.5.6). The atomic radii used in our program are as follows: C=1.70Å, N=1.55Å, O=1.52Å, S=1.75Å and H=1.09Å.

The structure in Fig. S3 is a peptide (min1.pdb) downloaded from an Amber tutorials website. The MSMS SES surface is generated using a probe radius=1.2Å, density=100 and high density=100, the highest values allowed for the two densities by MSMS as implemented in PMV. In MSMS these two densities determine the resolution for surface triangulation. MSMS restarts SES computation once with modifications to the atomic radii of the following three atoms: 66, 69 and 97. The SESs by our algorithm and by MSMS are very similar visually. Large differences appear only on regions with probe-probe interactions.

Figure S4 depicts the binding cavity of a 138-residue protein (human CRABPII) with a total of 2,098 atoms including protons. MSMS clashes with a *R_p_ =* 1.4&#x00C5;, density=100 and high density=100 though it could generate the surface with the same densities but a *R*_*p*_ &#x2014; 1.3&#x00C5; and with several restarts. Though the SES surfaces for the binding cavity by the two algorithms resemble each other greatly MSMS lacks the sharp-edges that demarcate different concave spherical polygons on intersected probes.

Figure S5 shows the SESs by our algorithm and by MSMS for a 1,328-residue protein with a total of 21,792 atoms. For large-sized proteins our algorithm remains to be robust and the fine details on the computed SESs remain to be clearly visible even both atomic spheres and probe spheres are each represented by 10, 242 vertices. In the contrary for large-sized proteins MSMS requires lower resolution for surface triangulation and clashes frequently when a high density value is used. Though the SESs by our algorithm and by MSMS are rather similar in general, the MSMS surface does not have well-defined arcs in regions with probe-probe intersections and has less fine details even with the highest possible densities. In addition there are some artifacts in the MSMS surface likely due to either the inadequate treatment of either probe-probe interactions or limited numerical precision or the bugs either in surface triangulation or in PDB file processing.

^1^Abbreviations: SES, solvent-excluded surface; SA, solvent-accessible; 2D, two-dimensional; PDB, Protein Data Bank.

---

**Figure S1:**
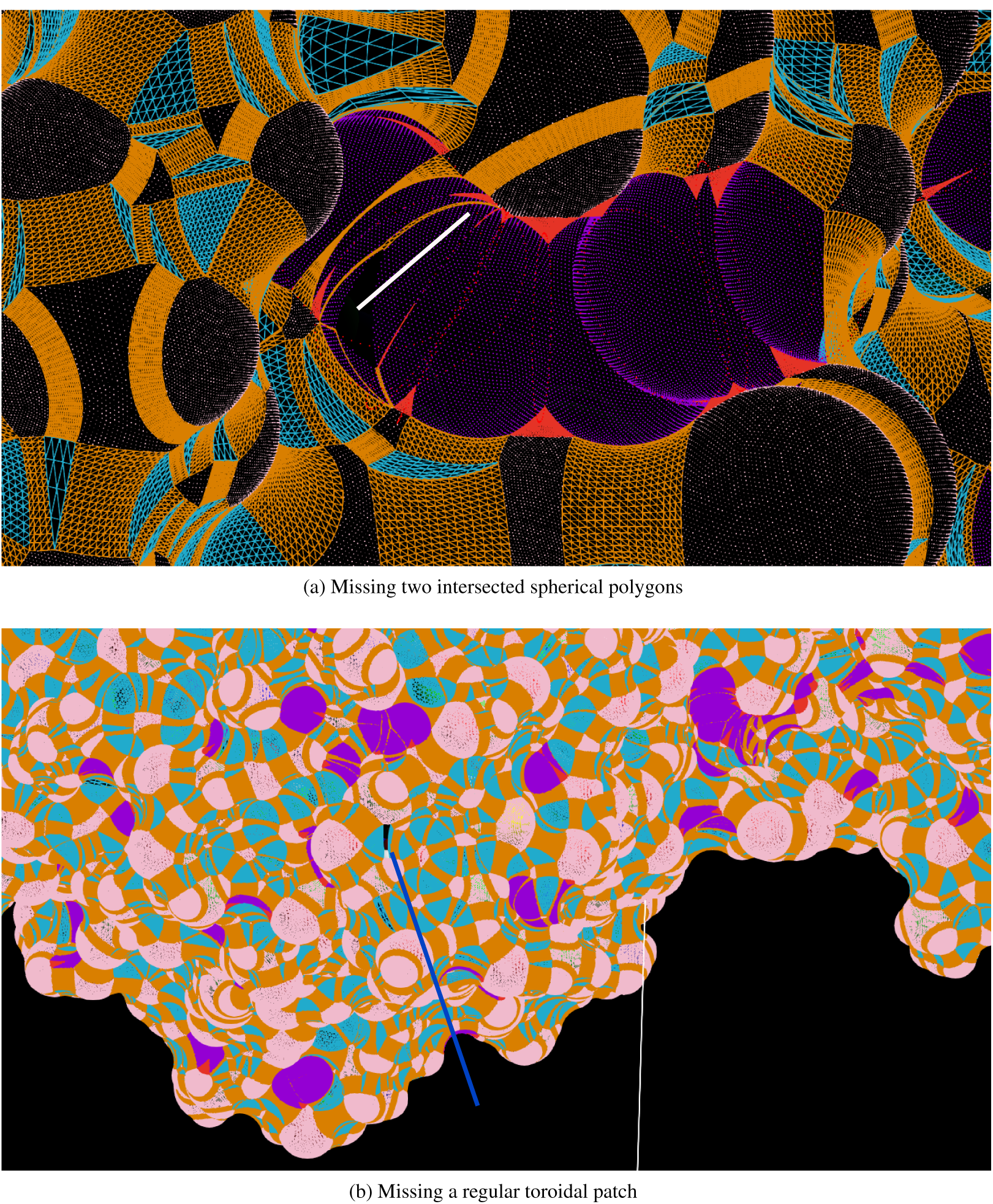
Two examples of missing patches. **(a)** A white line points to a SES region where two ∂_*p*_*s* on two intersected fixed probes are missing. The intersected ∂_*p*_*s* are rendered as points and colored in violet. The great circles of the intersected probes are rendered as points and colored in red. **(b)** A missing regular toroidal patch is indicated by a blue line. In both **(a)** and **(b)** the convex spherical polygons (*∂*_s_s) on accessible atomic spheres are rendered as points and colored in pink, the toroidal patches as wire-frames and colored in orange for regular tori and in red for spindle tori, and the *∂*_*p*_*s* on isolated probes are rendered as wire-frames and colored in cyan. The white line in **(b)** is an axis.

**Figure S2:**
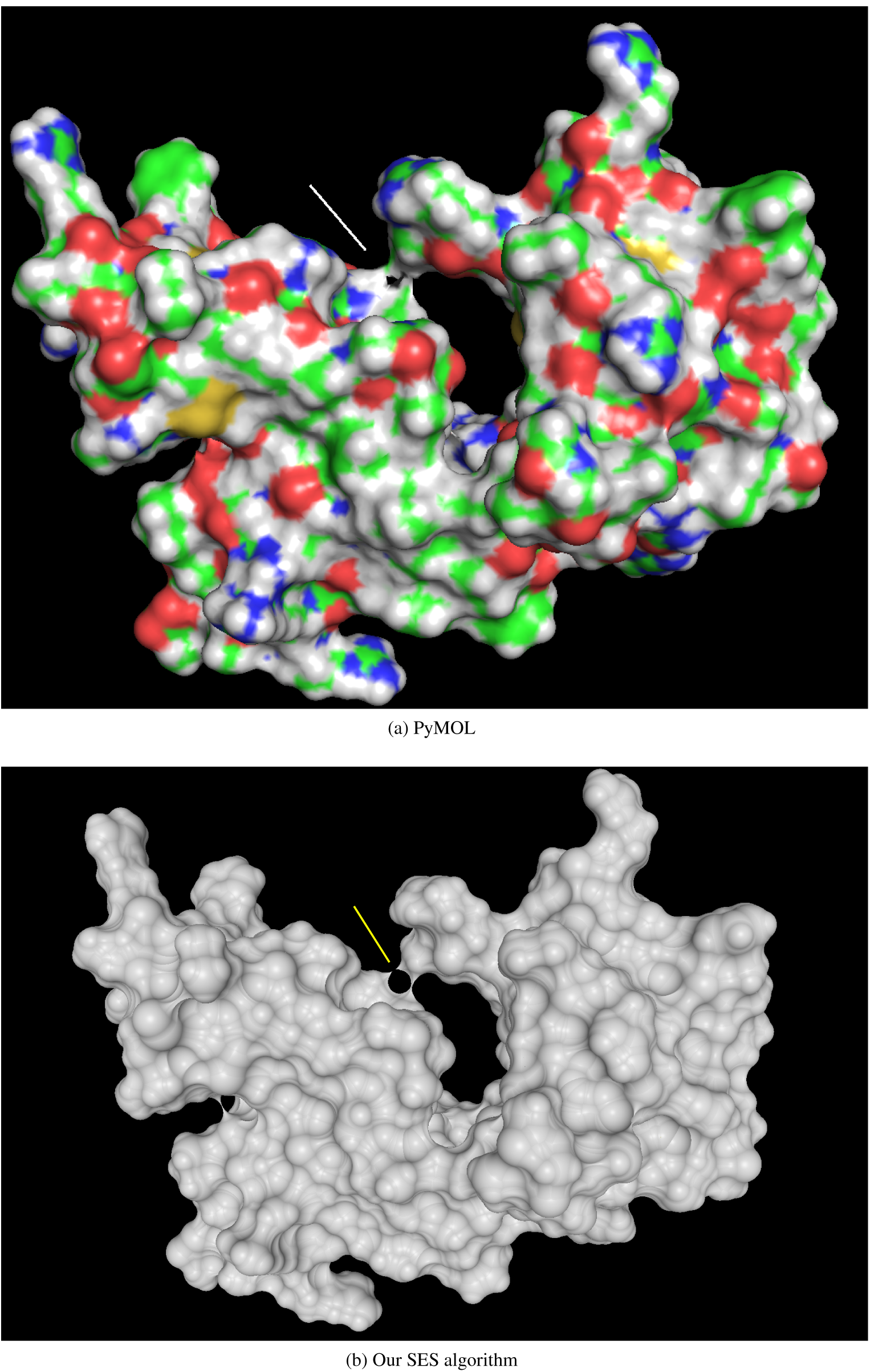
A defect in molecular surface by a grid-based algorithm. **(a)** The surface generated by PyMOL with a defect region indicated by a white line. **(b)** The SES generated by our algorithm for the same protein with the same region indicated by a yellow line. A classical Phong shading model with no specular light is used for lighting. A comparison between the two surfaces shows that the defects in PyMOL surface are caused by an inadequate treatment of probe-probe intersection. The protein is 1zmm (pdbid).

**Figure S3:**
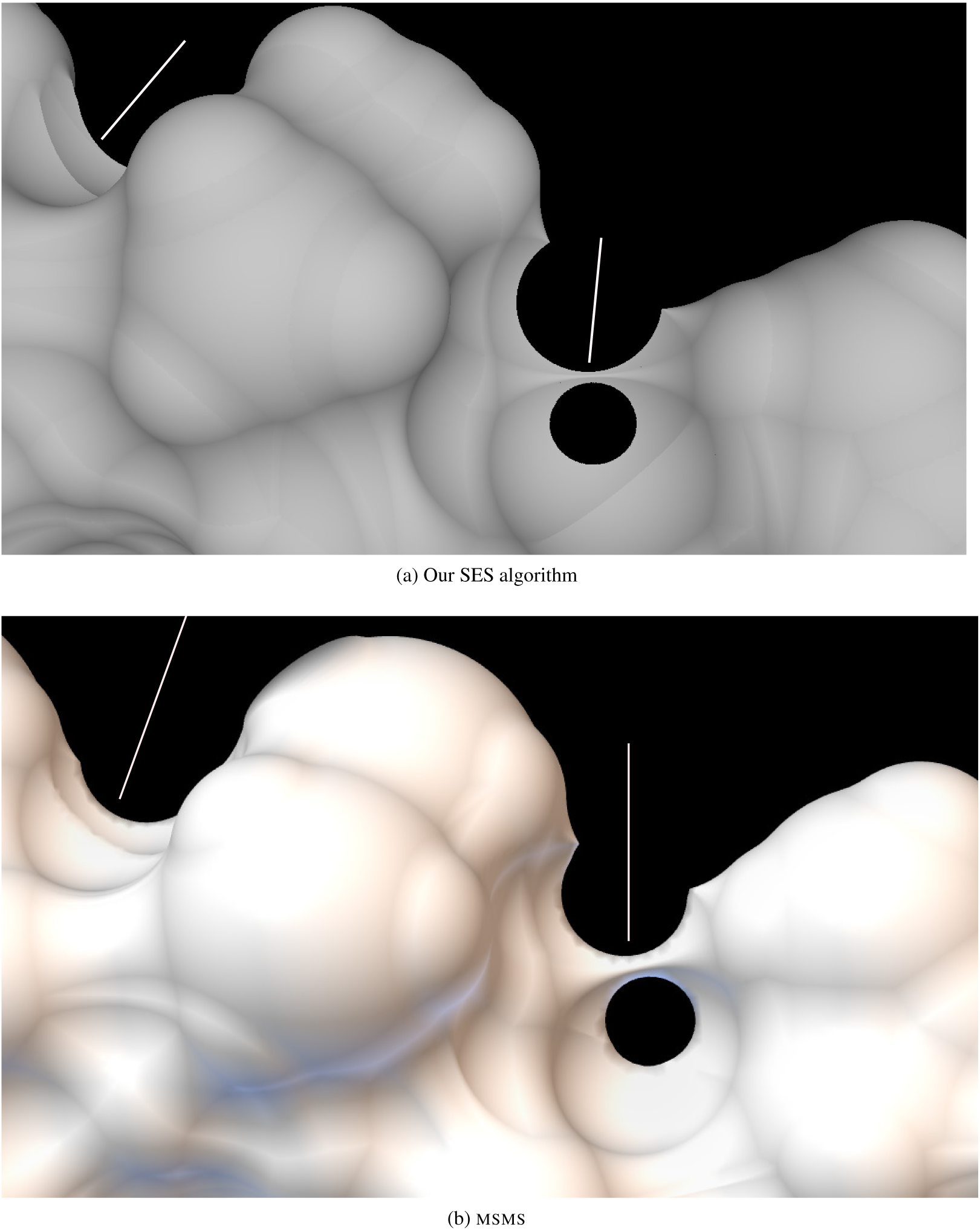
Intersected spherical polygons (*∂*_*p*_s) on fixed probes. The two regions with probe-probe intersections are indicated by two white lines. **(a)** In our algorithm the analytically-computed geometries (e.g. 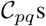) are used directly for surface triangulation. The SES is computed using 40, 962 vertices for sphere representation. **(b)** The same regions with probe-probe intersections are relatively rough in MSMS even with the highest densities. A classical Phong shading model with no specular light is used for lighting.

**Figure S4:**
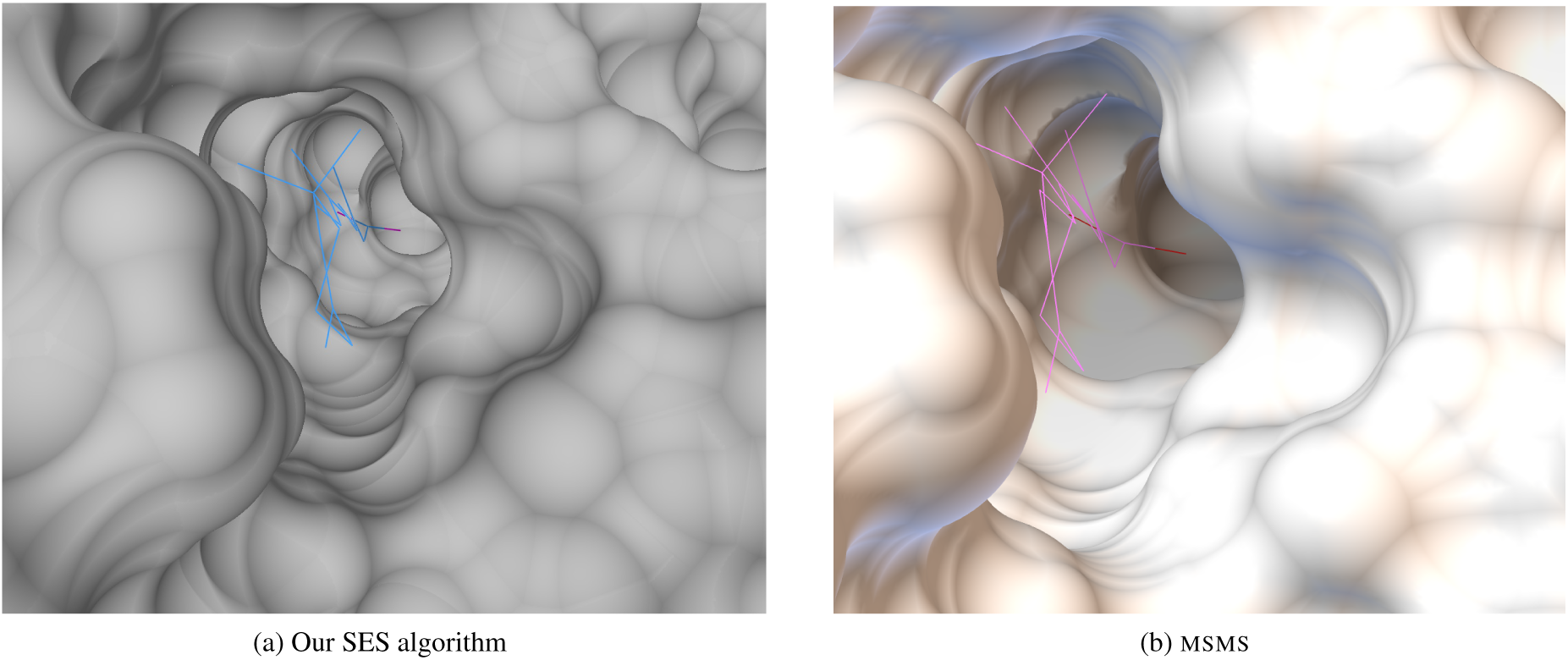
The binding cavity of human retinoic acid binding protein (type II). The **msms** surface is generated using a probe radius=1.3Å, density=100 and high density=100. The protein has 137 residues with a total of 2098 atoms including protons (pdbid: 2FR3). The bound ligand, an all*-trans-retinoic* acid, is depicted in line.

**Figure S5:**
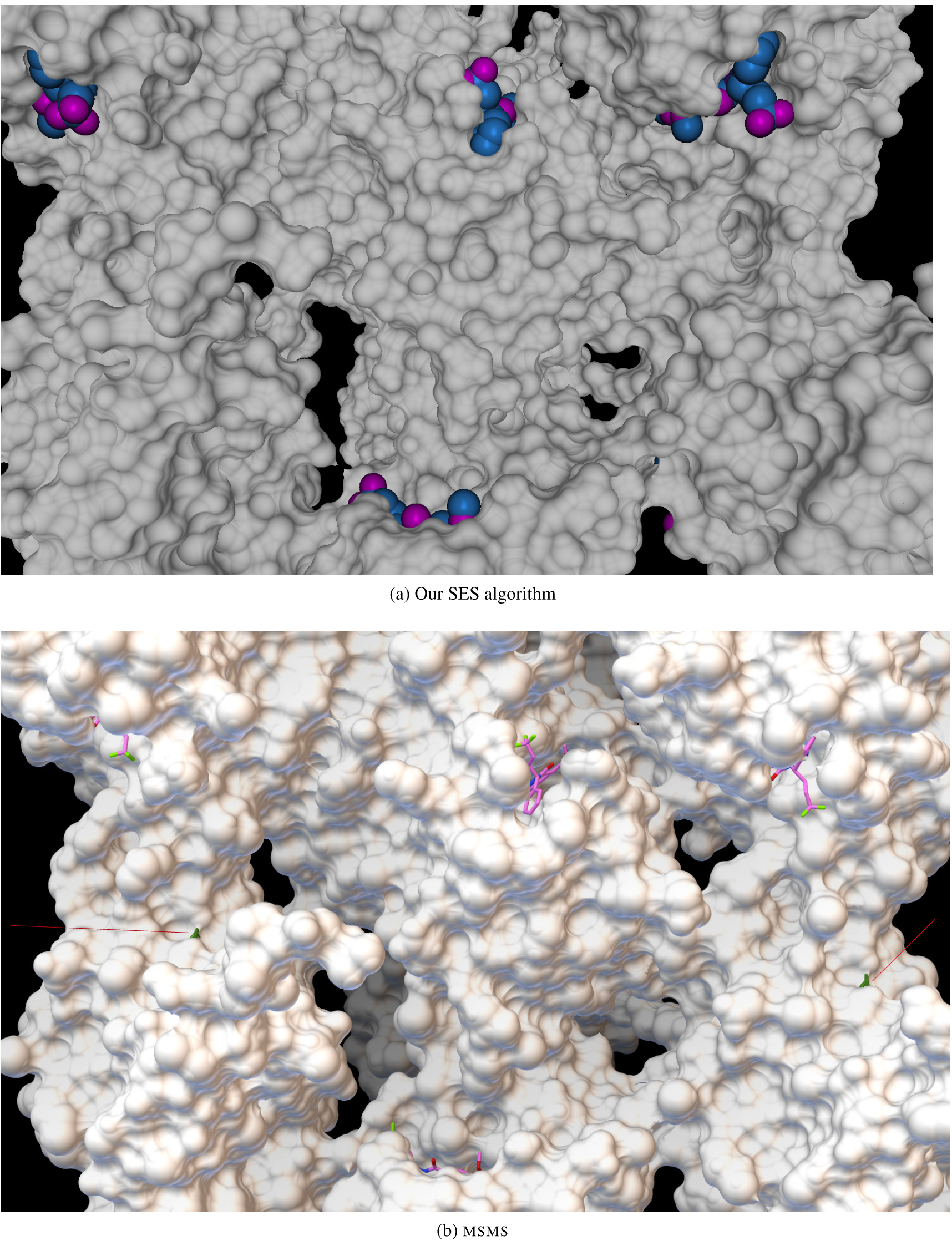
The SES of an oxidoreductase with inhibitors. It is a homo-octomer with each chain having 166 residues and a total of 21,792 atoms including protons (pdbid: 4ELF). **(a)** The SES surface is computed with 10, 242 vertices for sphere representation. **(b)** The MSMS surface is generated using a probe radius=1.3Å, density=20.0 and high density=40.0. There are artifacts in the MSMS surface as indicated by two red lines.^2^The case where an atom is surrounded by a single *∂*_*t*_(*i, j*) is treated differently. For ease of exposition we will make no more mention of it in the rest of the paper.

## Footnotes

1 Abbreviations: SES, solvent-excluded surface; SA, solvent-accessible; 2D, two-dimensional; PDB, Protein Data Bank.

2 The case where an atom is surrounded by a single *∂*_*t*_(*i; j*) is treated differently. For ease of exposition we will make no more mention of it in the rest of the paper.

